# Effect of lipopolysaccharide (LPS) on HAEC cells. Does nicotinamide N-methyltransferase sensitize HAEC cells to LPS?

**DOI:** 10.1101/2021.12.29.474421

**Authors:** Oksana Stępińska, Dorota Dymkowska, Łukasz Mateuszuk, Krzysztof Zabłocki

## Abstract

Treatment of endothelial cells with bacterial lipopolysaccharide (LPS) evokes a number of metabolic and functional consequences which built a multifaceted physiological response of endothelium to bacterial infection. Here effects of LPS on human aortic endothelial cells (HAEC) have been investigated. Among the spectrum of biochemical changes substantially elevated N-nicotinamide methyltransferase (NNMT) protein level was particularly intriguing. This important enzyme may potentially affect cellular metabolism by two means: direct regulation of methylnicotinamide level and availability of nicotinamide, that at least potentially may influence NAD^+^ synthesis, and regulation of S-adenosylmethionine concentration and therefore controlling methylation of many proteins including chromatin. This may have epigenetic consequences. This paper is focused on NNMT, despite the fact that in the presence of LPS additional effects of this compound mask pure (canonical) consequences of the elevated NNMT protein which are an increased MNA synthesis or reduced NAD^+^ level. On the other hand, however, it has been shown that silencing of the NNMT-encoding gene prevents several changes which are observed in control HAECs treated with LPS. They include significantly increased calcium response to thapsigargin (store-operated calcium entry), altered energy metabolism which is switched to anaerobic glycolysis and rearrangement of the mitochondrial network. However, a biochemical mechanism behind the protective consequences of the NNMT deficiency in cells treated with LPS remains unexplained.

## Introduction

Endothelial cells are directly exposed to many physiological and pathological factors transported in the blood. Bacterial lipopolysaccharides (LPS) belongs to the latter category. They form a family of large molecules containing three structural elements: a core oligosaccharide, an O-antigen, and a lipid A component [Bertani and Ruiz, 2018; Farhana and Khan, 2021; Whitfield et al., 2020]. LPS is an endotoxin produced by gram-negative bacteria which is neutralized under normal conditions by the innate immune system. This involves activation of intracellular signalling pathways which are initiated by activation of TLR4 receptors, evolutionarily conserved proteins which are responsible for recognizing bacteria-derived infections and form the first line of defending them. High LPS levels have certain toxic effects on cells, whereas low LPS levels promote cell proliferation. In sepsis cellular response to LPS is enormously strong and results in a generalized progressive inflammatory response [for rev. Fock and Parnova, 2021]. Epidemiological studies have indicated that LPS constitute a risk factor for diseases such as atherosclerosis and diabetes [Chao et al., 2017]. Inflammatory and immunological responses of endothelial cells stimulated with LPS have been broadly investigated. It has been found that LPS not only activates endothelial cells indirectly through inflammatory mediators, such as tumour necrosis factor α (TNFα), interleukin-1β (IL-1β), interferons (IFNs) and other released from macrophages and immune cells [Gabarin et al., 2021; Meng and Lowell, 1997] but also directly due to stimulation of NO generation and therefore supporting vasodilatation [Chang et al., 2000]. Moreover, LPS increases endothelial permeability and leads to endothelial barrier dysfunction [Schlegel et al., 2009]. Intensive studies on endothelium were mostly focused on a development of new therapeutic strategies as endothelial impairments are behind cardiovascular diseases and are a cause of excessive mortality in western countries in particular. On the other hand, cellular bioenergetics and the regulation of intermediary metabolic processes in endothelial cells are still unclear. It is commonly accepted that endothelial energy metabolism relies mostly on glycolytic ADP phosphorylation while the major mitochondrial role is to regulate intracellular calcium signalling and reactive oxygen species formation [Szewczyk et al., 2015]. However, in pathological conditions, metabolic relations and the contribution of particular processes to global cellular metabolism may be changed. Investigating biochemical mechanisms responsible for endothelial response to various stimuli may be helpful to a better understanding of pathological processes and finding new therapeutic approaches.

Our previously published data convincingly show that challenging endothelial cells with TNFα or palmitate affects mitochondrial network architecture and stimulates mitochondrial biogenesis which was interpreted as a pro-survival response to the stimuli applied at a relatively low concentration insufficient to induction of acute cell death [Drabarek et al., 2012; Dymkowska et al., 2017]. Furthermore, we found an incubation of endothelial cells with statins results in cellular stress response which was accompanied by an elevation of nicotinamide-N-methyl transferase (NNMT) [Dymkowska et al., 2021]. This enzyme catalyzes methyl group transfer from S-adenosylmethionine (SAM) to nicotinamide (NA). Reduction of NNMT gene expression was found to increase the detrimental effect of menadione. In turn, activation of NNMT/SIRT1 pathways seems to have a protective effect increasing the survival rate of endothelial cells in oxidative stress [Campagna et al. 2021]. Methylnicotinamide (MNA) is a modulator of several physiological functions of endothelium because of its anti-inflammatory and anti-thrombotic properties [Bartus et al., 2008; Chlopicki et al., 2007; Mateuszuk et al. 2020]. MNA was also found to have vasodilatory properties as it stimulates NO formation. Moreover, it has a multifaceted positive effect on lipid metabolism and storage, as well as lipoprotein and PGI2 levels [Nejabati et al., 2018]. Manifold effects of MNA on endothelial cells has recently been described in a comprehensive review [Begum et al. 2021].

Methylation of NA might potentially reduce its availability for NAD^+^ synthesis *via* the salvage pathway and therefore affect cellular redox processes. In turn, reduced accessibility of NAD^+^ could limit sirtuins-catalyzed deacetylation of many substrates including acetylated histones. Furthermore, an excessive NNMT activation may reduce SAM availability for other reactions and influence various methylation processes. Both events may have epigenetic consequences and influence gene expression due to affected chromatin modifications. Thus NNMT forms a junction between direct acute metabolic regulation and modified gene expression. Interestingly, overexpression of the NNMT gene in SH-SY5Y neuroblastoma cells significantly increased the expression of sirtuins 1, 2 and 3 encoding genes and a role of sirtuins 3 as a protein mediating effect of NNMT on the respiratory complex I was postulated. NNMT is expressed in a variety of tissues and organs and an elevated amount of this enzyme was found in many pathologies. Among them are Parkinson’s disease (neurons), cirrhosis (liver), pulmonary disorders and a variety of cancers [Roessler et al., 2005; Tomida et al., 2008; Sternak et al., 2010; Kim et al., 2010; Emanuelli et al., 2010; for rev see Roberti et al., 2021].

In view of all the aforementioned facts changes in cellular NNMT content and activity may be triggered in diverse pathological conditions and have very complex metabolic and physiological consequences concerning a broad spectrum of cellular functions. Therefore, the holistic depiction of the role of NNMT seems to be very puzzling. Our previous experiments on the energy metabolism of vascular endothelium were done with the use of immortalised human endothelial cells, EA.hy926. Here we have switched our interest to the primary human aortic endothelial cells (HAEC) and their metabolic response to bacterial lipopolysaccharide. We have focused our attention on the effects of LPS on mitochondria, intermediary energy metabolism and the putative role of NNMT in the cellular response to this lipopolysaccharide. We have found that incubation of HAEC cells with LPS substantially increases NNMT protein level and silencing of the NNMT-encoding gene modifies and mostly alleviates some other cellular changes which emerged as a response to the LPS treatment. However, we have not confirmed an expected direct link between an elevated NNMT level and cellular NAD^+^ content. Thus, in this study, we have pointed out the role of NNMT in the endothelial response to LPS while molecular mechanisms behind these effects remain obscure. A complete understanding of these effects needs further investigation.

## Material and Methods

### Cell culture and treatment

Human Aortic Endothelial Cells (HAEC) derived from two the same age male donors were purchased from Lonza. Cells were grown in Endothelial Cell Growth Medium BulletKit®-2 (EGM-2 BulletKit, Lonza) at 37°C in an atmosphere of 5% CO_2_ and 95% air. The culture medium was supplemented with conveniently packaged single-use aliquots called SingleQuots, containing: human recombinant epidermal growth factor, human fibroblast growth factor, vascular endothelial growth factor, ascorbic acid, hydrocortisone, human recombinant insulin-like growth factor, heparin, 2% foetal bovine serum and gentamicin with amphotericin (Lonza). Cells were passaged every three days. Confluent cells were exposed to 100 ng/ml LPS (Escherichia coli O111:B4, List Biological Laboratories, #421) for a period between 0.5 h to 24 h, as indicated in figure legends. In all experiments, cells of the third passage were used.

### NNMT gene silencing

The cells were cultured for 24 hours before starting the silencing procedure. A freshly prepared mixture of NNMT specific siRNA (Ambion, ThermoFisher Scientific, #4390826) and jetPrime transfection reagent (PolyPlus transfection) was added to the confluent cells for 24 hours. Then the silencing solution was replaced with a fresh culture medium and incubated for another 24 hours before further treatment with LPS. The scrambled siRNA (Ambion, ThermoFisher Scientific, #4390847) was used as a control. The efficiency of NNMT gene silencing was determined by the Western Blot technique.

### Cell lysis, and Western Blot analysis

Cell lysates were prepared as previously described [Drabarek et al., 2012; Dymkowska et al., 2014]. Proteins were separated by polyacrylamide gel electrophoresis (PAGE) under denaturing conditions in the presence of 0.1% sodium dodecyl sulphate (SDS, BioShop) [Laemmli, 1970]. After transferring to the PVDF membrane (Millipore) NNMT protein was detected with the use of a specific primary antibody (Santa Cruz Biotechnology, sc-376048); anti-rabbit and anti-mouse secondary antibodies conjugated with horseradish peroxidase (HRP) were obtained from Abcam. For HRP detection (Fusion FX, Vilber Lourmat) chemiluminescent substrate Immobilon Classico (Merck Millipore) was used. The optical density of bands corresponding to defined proteins was determined densitometrically with the BIO-1D software (Vilber Lourmat) and expressed in relation to β-actin used as a loading control (Sigma, A-3854).

### Immunocytochemistry

For visualization of the mitochondrial network architecture, confluent cells grown on the collagen-coated coverslips (ϕ12 mm) were loaded with 100 nM MitoTracker Red CMXRos (Molecular Probes) as previously described [Dymkowska et al., 2021]. After gentle rinsing, the cells were fixed with 4% paraformaldehyde, rinsed with PBS supplemented with 5% BSA, permeabilized with 0.1% Triton X-100 in 5% BSA and rinsed overnight with 1% BSA containing PBS at 4°C. Finally cells were stained with Actin-Stain 488 Phalloidin (1:1000; Cytoskeleton) and with 2 µg/ml Hoechst 33342 (ThermoFisher Scientific) Finally, cells were sealed in VECTASHIELD Mounting Medium (VECTOR Laboratories). Fluorescence microscopy analysis was carried out using a Zeiss Spinning Disc microscope.

### Metabolite determination using the HPLC method

To measure glycolytic intermediates, TCA metabolites and adenine nucleotide intracellular content the cells were rinsed with cold 0.3 M mannitol. Then cells monolayers were extracted with the mixture composed of methanol: acetonitrile: water (2:2:1). The extracts were collected and centrifuged at 4 °C. The supernatants were stored at -80 °C for further procedures. Pellets were stored for an amount of total protein measurement. Before metabolite assay to each sample 140 pmoles of the appropriate standard was added and then selected metabolites were separated and quantitively analyzed with the use of an ion-exchange chromatography system (Dionex ICS-3000 chromatograph, Thermo Fisher Scientific Inc.) coupled with Waters ZQ mass spectrometer (Waters Corporation). Metabolite separation was performed on AS11-HC high-capacity anion-exchange column using 1-80 mM KOH gradient as the solvent. In the case of ATP, ADP and AMP measurements, an integrated with Dionex chromatograph UV-detector (260 nm) was used. All samples were measured three times and results were quantified both to the reference standard and amount of cell protein and presented as a mean of replicates.

To determine the extracellular methyl nicotinamide (MNA) level 100 µl of culture medium was collected from each dish and frozen at -80°C. Cell monolayers were rinsed with PBS, trypsinized and centrifuged. The pellets were stored for an amount of total protein measurement. Aliquots of 50 μl of defrosted medium were spiked with 5 μl of internal standard (deuterated analogues of the analyte) at the concentration of 25 μg/ml. Samples were subjected to deproteinization with 100 μl of acetonitrile acidified with 0.1% formic acid), vortexed, cooled at 4°C for 15 min and centrifuged (15 000 x g, 15 min, 4°C). Supernatants were injected into the LC column. Chromatographic analysis was performed using an UltiMate 3000 LC system (Thermo Scientific Dionex, Sunnyvale, CA) consisting of a pump (DGP 3600RS), a column compartment (TCC 3000RS), an autosampler (WPS-3000TRS), and an SRD-3600 solvent rack (degasser). Chromatographic separation was carried out on an Aquasil C18 analytical column (4.6 mm x 150 mm, 5 mm; Thermo Scientific). MNA was eluted with the mobile phase consisting of acetonitrile (A) and 5 mM ammonium formate (B) in isocratic elution (80:20 v/v) at the flow rate of 0.8 ml/min. MNA detection was performed with the use of a TSQ Quantum Ultra mass spectrometer equipped with a heated electrospray ionization interface (HESI-II Probe) (Thermo Scientific, Waltham, MA, US). The mass spectrometer was operating in the positive ionisation using selected reactions monitoring mode (SRM), monitoring the transition of the protonated molecular ions (for MNA Precursor [m/z] = 137, Product [m/z] = 94). Data acquisition and processing were accomplished using Xcalibur 2.1 software.

To measure cellular NAD^+^ content the cells were rinsed with cold PBS and then extracted with an ice-cold 10% solution of perchloric acid. Then the cells were harvested, transferred to Eppendorf tubes, forced through the thin needle, vortexed and incubated on ice for 15 minutes. Then, lysates were centrifuged at 15 000 x g at 4°C for 5 min. The supernatants were carefully transferred to fresh tubes and neutralized with 3 M potassium carbonate and centrifuged again at 15 000 x g at 4°C for 5 min. Supernatants were transferred to fresh tubes and stored at -80°C. NAD^+^ measurements were made according to the method described by Yoshino and Imai [2013].

### Calcium measurement

Cytosolic Ca^2+^ concentration was measured with the fluorescent probe Fura-2-AM. Cells grown on coverslips previously coated with collagen I (Sigma) were loaded with 2 μM Fura-2-AM in the culture medium for 30 min at 37°C in darkness. The cells were then washed twice with the solution composed of 5 mM KCl, 1 mM MgCl_2_, 0.5 mM Na_2_HPO_4_, 25 mM HEPES, 130 mM NaCl, 1 mM pyruvate, 5 mM D-glucose, and 0.1 mM CaCl_2_, pH 7.4 and the coverslips were mounted in a cuvette containing 3 ml of either the nominally Ca^2+^-free assay solution (as above but 0.1 mM CaCl_2_ was replaced by 0.05 mM EGTA). The fluorescence was measured at room temperature in a Hitachi F-7000 fluorimeter set in the ratio mode at 340 nm/380 nm excitation and 510 nm emission wavelengths. At the end of each experiment the Fura 2 fluorescence was calibrated by the addition of 8 μM ionomycin to determine maximal fluorescence followed by the addition of EGTA to complete removal of Ca^2+^ [Zabłocki et al. 2005].

### Oxygen consumption

HAEC cells were grown in Seahorse XFe96 polystyrene tissue culture plates (SeahorseBioscience Europe) and stimulated with LPS upon reaching confluence. Before measurement, cells were incubated in Seahorse XF DMEM assay medium (without phenol red and Sodium Bicarbonate) containing 10 mM glucose, 1 mM pyruvate and 2 mM glutamine in a non-CO_2_ incubator at 37 °C for 1 h. Oxygen consumption rate (OCR) was measured every 3 min with mixing of 3 min in each cycle, with 4 cycles per step using Seahorse XFe96 Analyzer (Agilent). Cell Mito Stress Test was used for assessing mitochondrial function. The sequential addition of oligomycin A (1.5 µM), FCCP (1.0 µM), antimycin A/rotenone (0.5 µM) dissolved in DMEM assay medium allowed for the calculation of OCR linked to ATP production, maximal respiration capacity and spare respiratory capacity. Basal respiration was measured prior to injection of oligomycin A. Finally, the data were normalized according to the total protein content in each well.

### Expression of the results

The results are presented as means of ratios of treatment (experimental group) to control values ± SD for the number of separate experiments. Statistical significance of differences (p-values less than 0.05) was calculated using a one-way analysis of variance (ANOVA). Proper Multiple Range Tests was used for comparisons between experimental groups.

## Results

### Response of control HAECs to LPS

In all experiments, LPS was used at a concentration of 100 ng/ml. It was the lowest one that was found to induce cellular response expressed as an elevation of the adhesion molecules ICAM-1 and VCAM-1 and protein involved in stress response levels (SOD1 and 2, HO-1 and COX-2). As these results confirm previously published experimental data obtained with the use of various cells which are dispersed amongst a number of papers [Dayang et al., 2019; Menden et al;., 2013; Sampath et al., 2009] they are not shown here and considered as auxiliary information specifically focused on HAEC cells.

Fluorescent microscopy analysis of HAEC cells treated with lipopolysaccharide indicates transient changes in the architecture of the mitochondrial network (Fig. 1) Mitochondria evenly dispersed within control cells became more condensed in the perinuclear space in the 6^th^ hour after the addition of LPS into the growth medium. Interestingly, the architecture of mitochondrial network observed after an additional 18 h seems to be substantially restored.

**Fig. 1.**
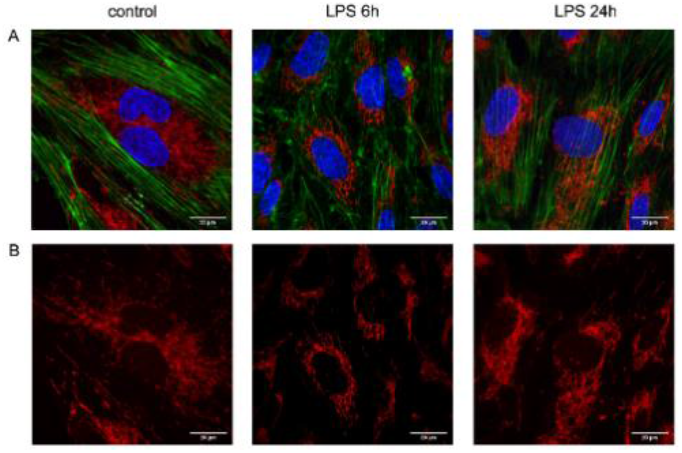
Effect of LPS on mitochondrial network organization. Green – actin cytoskeleton stained with Acti-stain 488 phalloidin, Red – mitochondria labelled with MitoTracker CMXRos, Blue – nucleus with Hoechst 33342.

LPS also strongly affects oxidative phosphorylation in HAEC cells but this effect in contrast to mitochondrial network organization does not reverse.

Fig. 2A shows representative traces obtained with the use of the Agilent Seahorse XFe96 analyzer. Although at this stage of the experiment the traces are not normalized to the cellular protein level they may be considered reliable as the cells were always seeded at the same density and the experiments were performed after the monolayer had reached complete confluency. Thus the number of cells in each well may be assumed as the same with a satisfying approximation. Fig. 2 B also points out changes in respiratory chain capability, which represents the fraction of the respiratory chain activity presumably attributed to covering energy demands for ADP phosphorylation. These effects are accompanied by deregulation of Ca^2+^ homeostasis that is expressed by an activated SOCE (Fig. 2C), and transiently increased ACC phosphorylation (Fig. 2E). The latter indicates an elevation of AMPK activity. In addition to these effects, LPS induces substantial changes in the proportion of cellular content of glycolytic and tricarboxylic acids cycle (TCA) intermediates (Fig. 3). Cross-over analysis clearly indicates transitory inhibition of pyruvate kinase followed by complete recovery and reversed ratio between PEP and pyruvate. (Fig. 3).

**Fig. 2.**
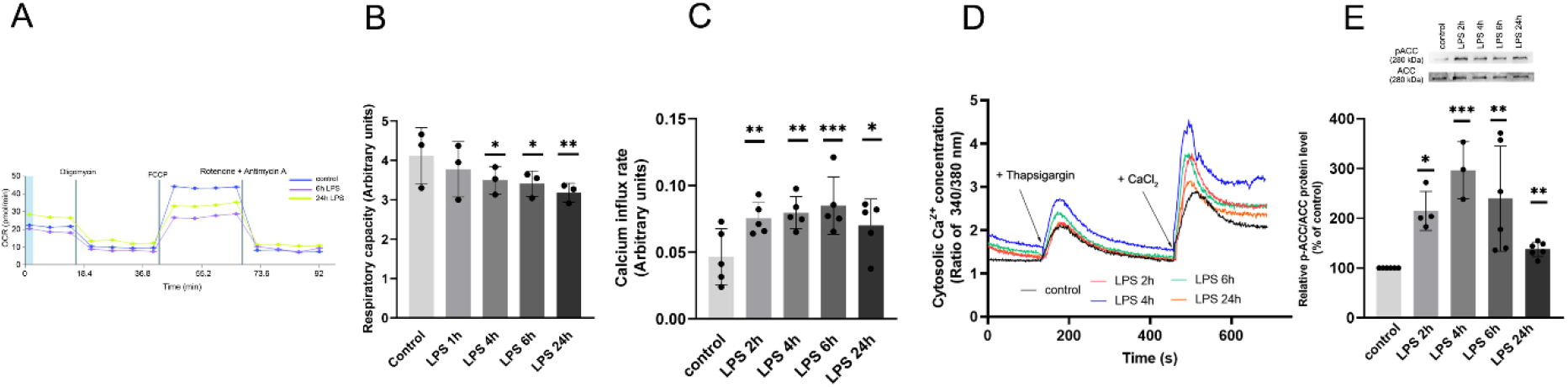
Effect of LPS on oxygen consumption, store-operated calcium entry and ACC phosphorylation (anti-p-ACC antibody, Cell Signaling, #3661) in HAECs. Row data obtained during direct oxygen consumption (representative experiment) and mitochondrial respiratory capacity of control and LPS-treated cells. Slopes after the addition of CaCl_2_ reflect rates of [Ca^2+^]c elevation due to activated store-operated calcium entry. Data show mean values ± SD for n = 3-5, *p<0.05, **p<0.005, ***p < 0.002.

**Fig. 3.**
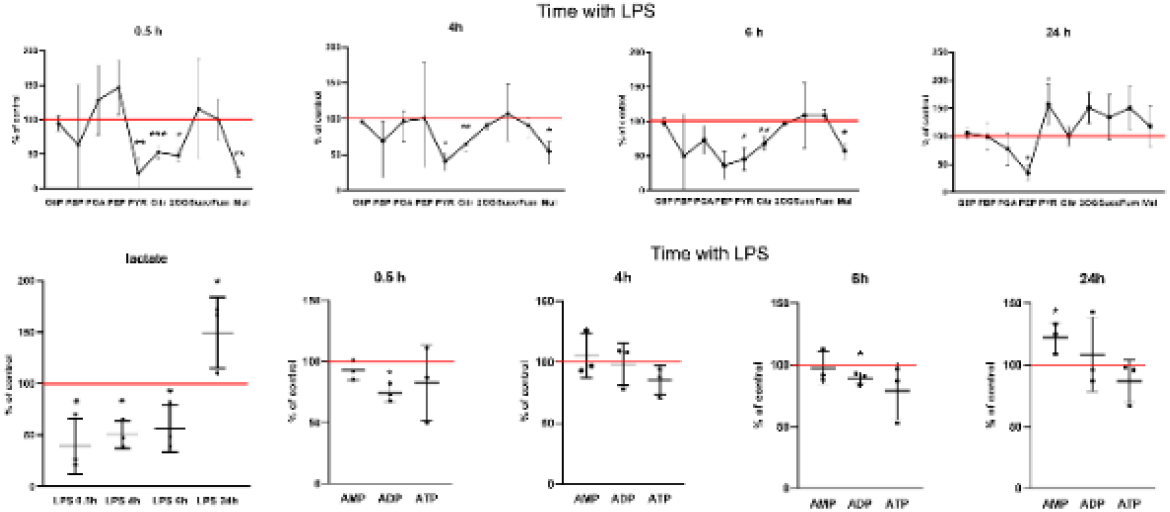
Effect of LPS on the relative metabolite and adenine nucleotide content. Cross-over graphs show mean values ± SD of the relative change of tested parameter vs. untreated control cells assumed as 100% (red line), n=3, *P<0.05, **p <0.005, ***p<0.0005.

These changes meet a substantial increase in lactate formation which follows a significant drop in its concentration shortly after the addition of LPS to the HAECs culture. Also, a huge cross-over effect between PEP and pyruvate apparently indicates a reduced flow between PEP and pyruvate at the beginning of treatment with LPS and subsequent reversal of this proportion suggests a relative elevation of pyruvate kinase activity. Pyruvate accumulation may also be caused by an inhibition of the further steps of its oxidation by pyruvate dehydrogenase and then in the TCA cycle. Excess of pyruvate is converted to lactate by lactate dehydrogenase. Effects of LPS on the adenine nucleotide content seem to be statistically insignificant though it exhibits the same tendency after 4, 6 and 24 h of the treatment. Thus some decrease in ATP level could be considered to be potentially relevant. Gradual reduction of the respiratory chain activity may also cause an excessive accumulation of TCA cycle intermediates. Finally, endothelial metabolism became more glycolytic but still efficient to deliver an ample amount of ATP. Our previously published data [Dymkowska et al., 2021], as well as results published by other authors [Akar et al., 2020], indicate that treatment of cells with various stimuli results in an elevation of NNMT content and this effect may be related to changes in cellular energy metabolism [Mistry et al., 2020]. Thus we also tested the putative effect of LPS on NNMT level in HAECs (Fig. 4).

**Fig. 4.**
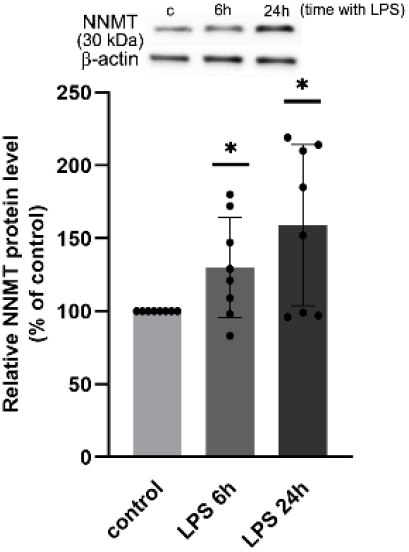
LPS increases NNMT level. Bars show relative data from 7 independent experiments. NNMT protein level was assumed to be 100%. Data shown as Mean ± SD, n = 7, * p <0.05. Above the bars, a Western Blot from one experiment is shown.

Despite these changes, the amount of MNA released from cells treated with LPS to the extracellular milieu is much lower than that in the case of untreated HAECs (Fig. 7. compare the first bars from each set of three). It shows that an LPS-induced increase in the amount of NNMT detected by the Western Blot approach does not correspond with NNMT activity at least estimated on basis of extracellular MNA level. Moreover, this partially transient descendant effect on MNA level is not accompanied by any changes in NAD^+^ content (Fig. 8, compare the first bars from each set of three). To test whether an LPS-induced increase in the NNMT level is of any importance for cellular metabolic response to this lipopolysaccharide or is unrelated to these effects, HAEC cells with silenced NNMT-encoding gene were used.

### Response of HAECs with silenced NNMT gene to LPS

An incubation of HAECs with silenced NNMT-encoding gene reduces NNMT protein level (Fig. 5A) and this effect persists for at least 72 hours, i.e. 24 h after addition of siRNA plus 24 h in the presence of LPS (Fig. 6). It is accompanied by a reduction in the amount of released MNA. However, decreased amount of methylated nicotinamide is not coupled with a significant elevation of NAD^+^ content (Fig. 5C).

**Fig. 5.**
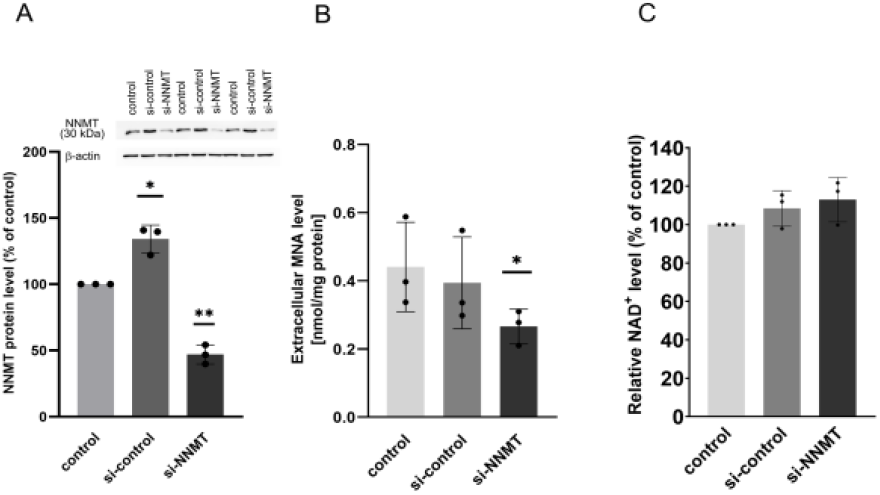
Effect of silencing of the NNMT-encoding gene and on NNMT protein level, MNA generation and NAD^+^ content 48 hours after addition of siRNA to the confluent cells. NNMT level was estimated with Western Blot. The representative result is shown above the graph. Data shown as mean±SD, n=3-8; *p<0.05, **p<0.005.

**Fig. 6.**
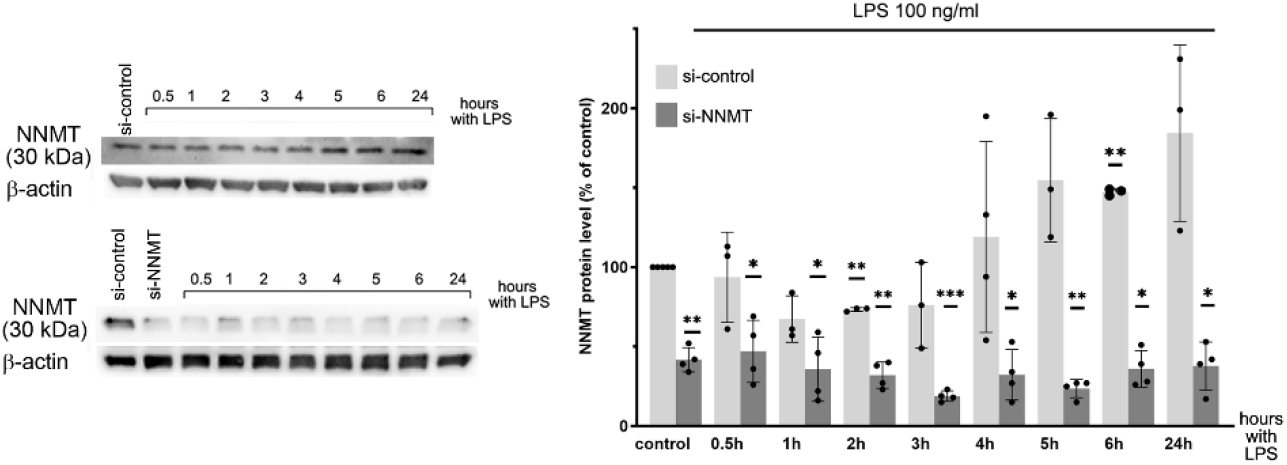
Effect of LPS on NNMT level in HAEC cells with silenced NNMT gene. Each bar indicates mean values ± SD of the relative change of tested parameter vs. untreated control cells (no LPS) assumed as 100%, n=3-8, *p<0.05; **p<0.005, ***p<0.0005. A representative Western Blot is shown on the left.

Furthermore, preincubation of cells with scrambled siRNA does not diminish the stimulatory effect of LPS on the NNMT level (Fig. 6). As expected, in cells with silenced NNMT gene LPS does not increase NNMT level.

Fig. 7 confirms that silencing the NNMT gene reduces the amount of extracellular MNA (the first set of three bars; similar but independently obtained result is shown in Fig. 5 B). LPS reversibly reduces MNA formation in control cells while in cells with silenced NNMT gene this inhibitory effect is much deeper and persist at least for 24 hours. Finally, cellular NAD^+^ content remains unchanged regardless of the mechanisms behind the reduced MNA formation (Fig. 8).

**Fig. 7.**
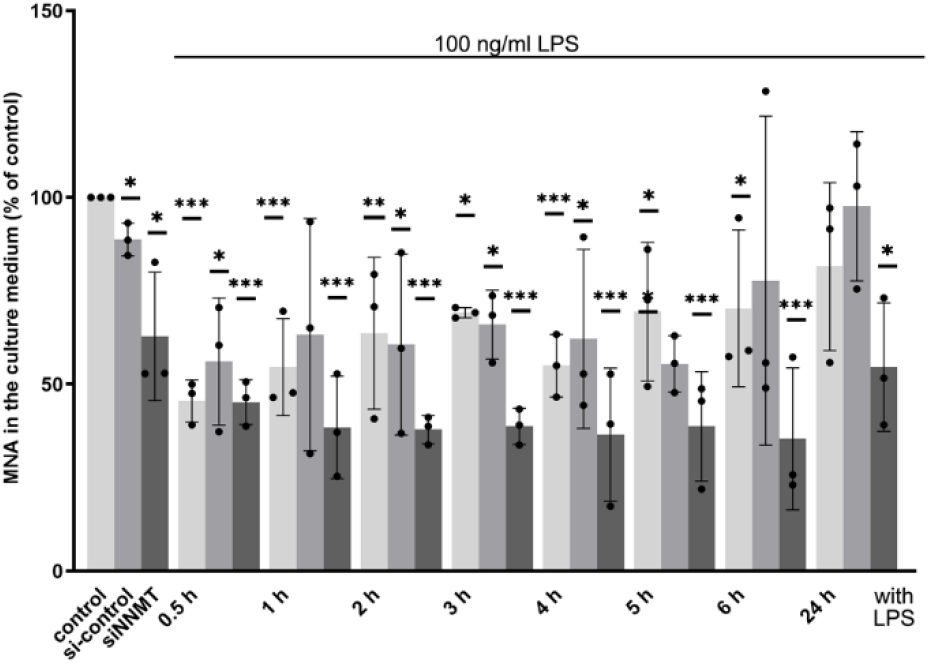
Effect of LPS on MNA content in HAEC cells with silenced NNMT gene. Each bar indicates mean values ± SD of the relative change of tested parameter vs. untreated control cells (no silencing, no LPS) assumed as 100%, n=3-8, *p<0.05; **p<0.005, ***p<0.0005.

**Fig. 8.**
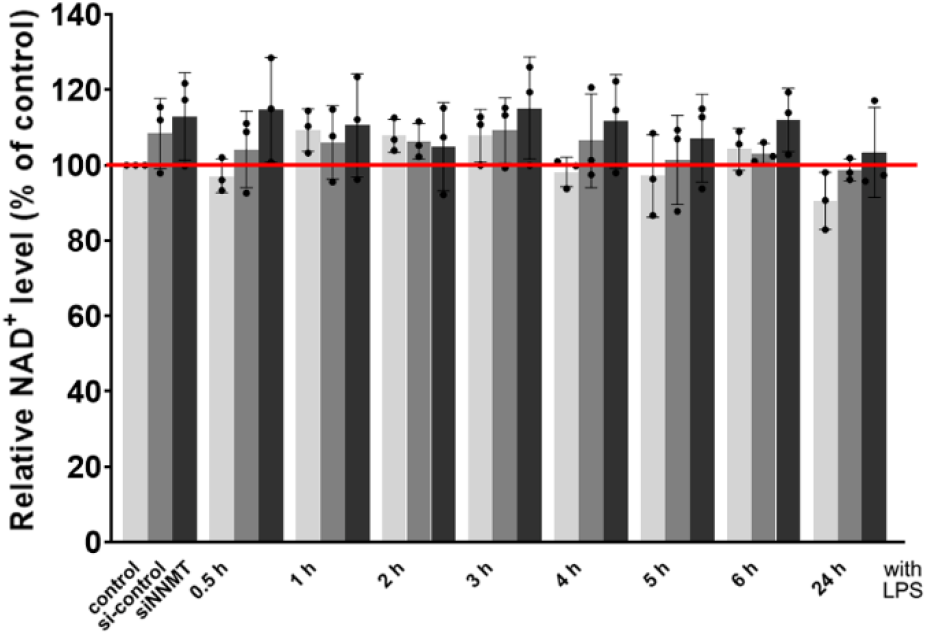
Effect of LPS on NAD^+^ content in HAEC cells with silenced NNMT gene. Each bar indicates mean values ± SD of the relative change of tested parameter vs. untreated control cells (no silencing, no LPS) assumed as 100%, n=3.

Fig. 9 shows that LPS-induced changes in the mitochondrial network architecture in HAECs cells are less visible in cells with silenced NNMT gene than in their equivalents treated with scrambled RNA. A comparison of these two populations indicates that silencing of the NNMT-encoding gene weakened the LPS-increased tendency of the mitochondrial network to be positioned more perinuclearly than in the case of control cells (see Fig. 1) or incubated with the scrambled RNA.

**Fig. 9.**
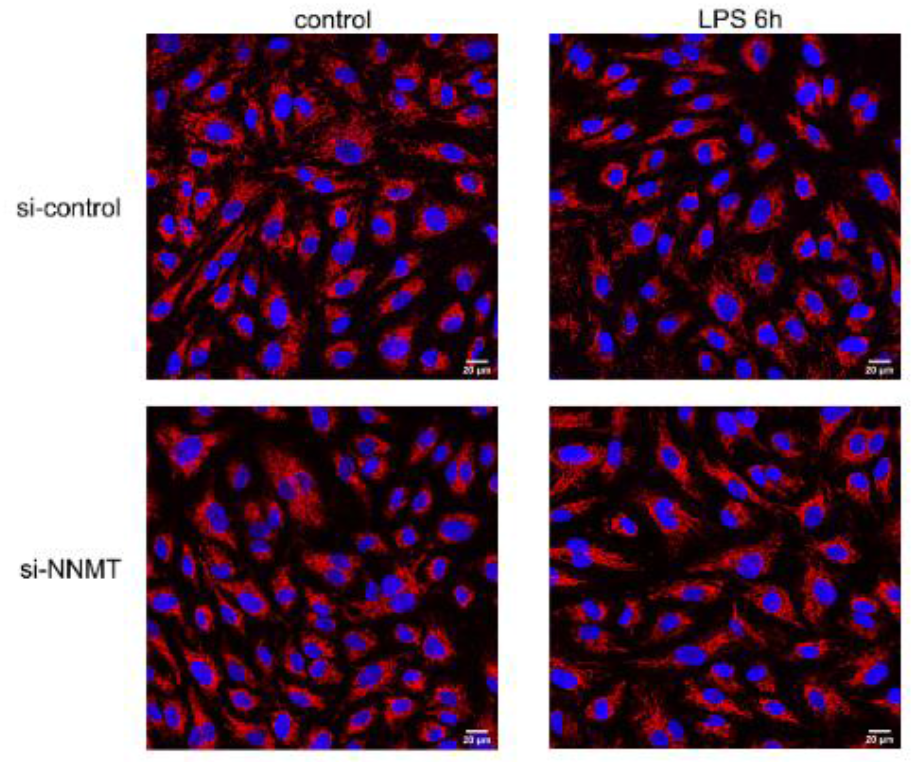
Effect of LPS on the mitochondrial network in HAEC cells with silenced NNMT-encoding gene. Red – mitochondria labelled with Mitotracker CMXRos, blue – nucleus with Hoechst 33342.

Fig. 10 shows that the LPS-evoked effect on the relative metabolite content in HAEC cells is visibly weakened in cells with silenced NNMT-encoding gene. As shown in Fig. 10 column A, LPS substantially reduces the level of most of the metabolites involved in glycolysis and the Krebs cycle in cells with unaffected NNMT gene expression (scrambled RNA) while this effect is less profound in cells with silenced NNMT (Fig. 10 column B). Fig. 10 C clearly shows that silencing of NNMT encoding gene diminishes or reverses effects of LPS visualised in Fig. 10 A. Since LPS does not affect the NAD^+^ level under any experimental conditions applied (Fig. 8), the availability of this dinucleotide does not seem to be a limiting factor that could control glycolysis and TCA. In other words silencing of the NNMT gene accelerates the recovery of metabolite profiles changed due to treatment with LPS but the biochemical mechanism beyond this effect is not clear. Results concerning metabolic profiles though fully convincing, should be interpreted cautiously as the procedure of cell transfection itself is a cause of some effects (compare Fig. 5 and Fig. 10 A). The effect of the NNMT-gene silencing itself on the metabolite profile of cells treated with scrambled RNA is shown in Fig. 10 D. Finally, silencing of the NNMT gene seems to gently prevent LPS-evoked stimulation of SOCE that was observed in cells with unaffected expression of NNMT (Fig. 11).

**Fig. 10.**
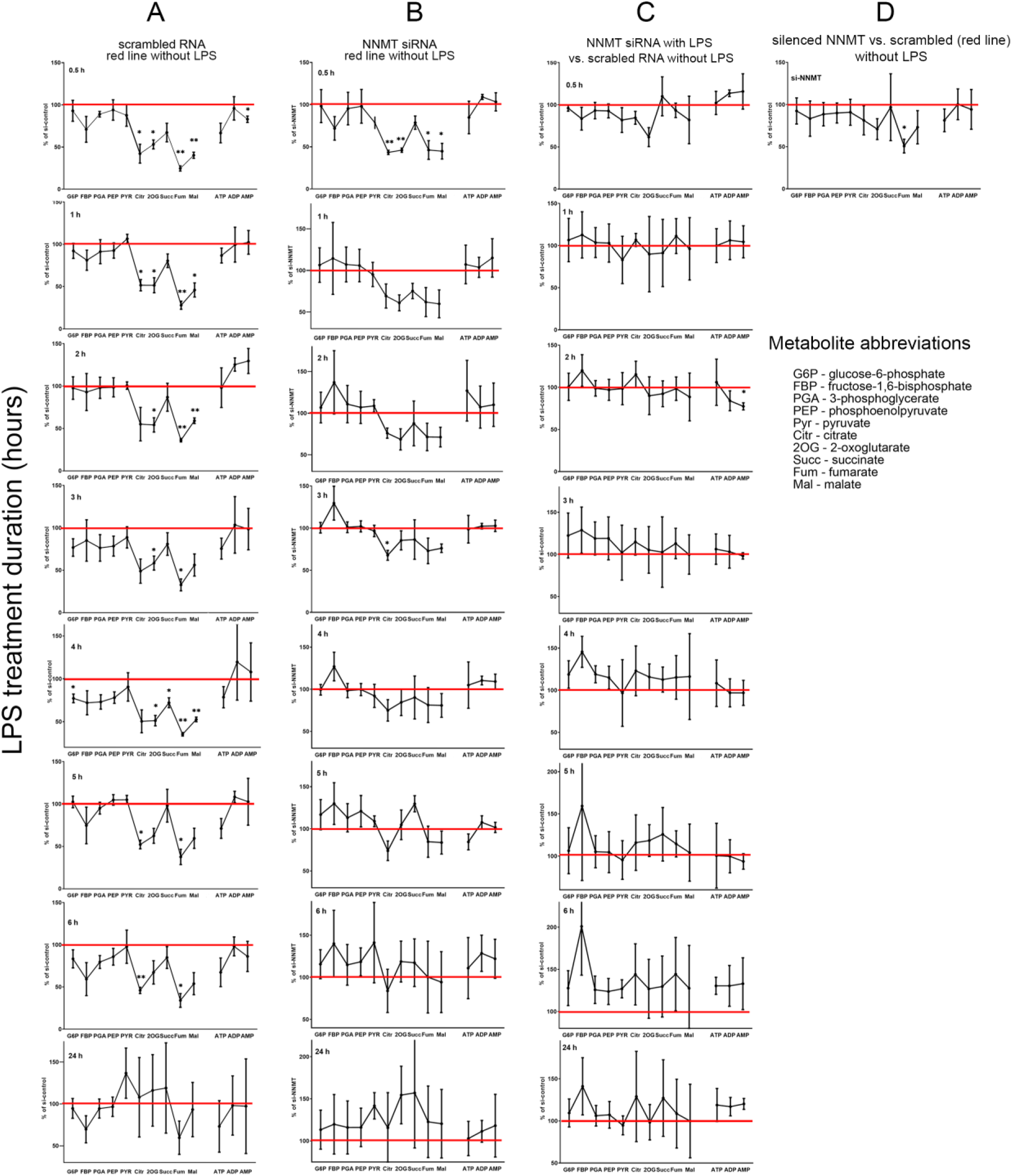
Effect of LPS and NNMT gene silencing on glycolytic and Krebs cycle metabolite content in HAEC cells. Column **A:** Effect of LPS on cells treated with scrambled RNA vs. without LPS; Column **B**: Effect of LPS on cells treated with siRNA (silenced NNMT gene) vs. without LPS. Column **C**: Effect of NNMT gene silencing plus LPS vs. cells treated with scrambled RNA without LPS, n = 3, *p<0.05, **p<0.005.

**Fig. 11.**
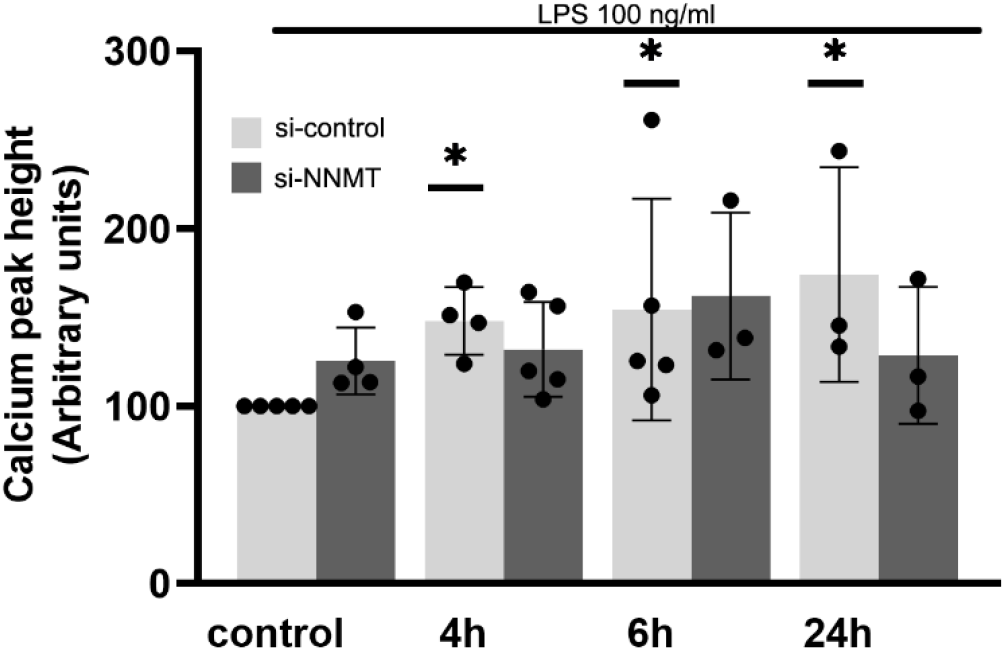
Effect of LPS on thapsigargin-induced calcium response in HAEC cells treated with scrambled RNA and after NNMT gene silencing, n=3-5, *p<0.05.

## Discussion

Effects of LPS on endothelial physiology, inflammatory response and NO generation have been broadly discussed for a long time[Chao et al., 2017; Dauphinee et al., 2006; Dayang et al., 2019; Li et al., 2017; You et al., 2021]. However, a detailed understanding of biochemical processes particularly focused on energy metabolism in human endothelium upon treatment with LPS needs further study. This issue seems to be particularly interesting in view of the fact that bacterial lipopolysaccharide substantially affects endothelial signal transduction which is behind the physiological responses of endothelial cells to a variety of stimuli. Calcium signalling and energy metabolism are mutually dependent factors that are of high importance for all cell types however, endothelial ones have been relatively poorly characterized in this matter. LPS was found to induce FADD-mediated endothelial cells apoptosis [Choi et al., 1998] and many studies are focused on the harmful effects of lipopolysaccharide in endothelial cells leading to reduced viability[Chen and Song, 2020]. Moreover, LPS-induced and Toll-like 4 receptor-mediated signalling in endothelial cells has been broadly investigated [Dauphinee and Karsan, 2006]. However, experimental efforts were primarily focused on searching for potential therapeutic approaches to prevent sepsis and serious cardiovascular complications.

In the present study, we have investigated the effects of relatively gentle treatment of cells with lipopolysaccharide to follow adaptive response without affecting cell viability. Thus, to minimize potential side effects of LPS, in all experiments presented here this stimulus was used at the lowest concentration and applied for the period which were established experimentally to be sufficient to elevate the protein level of adhesive molecules but not reduce cell survival.

### LPS affects endothelial bioenergetics

Mitochondria in endothelial cells are not in the centre of ATP generation but their role as a hub of cellular calcium signalling in all mammalian cells is unquestionable [Szewczyk et al., 2015]. An increased level of proteins involved in an oxidative stress response (preliminary data, not shown) together with substantially but reversibly changed architecture of the mitochondrial network indicate mitochondrial involvement in the response of HAECs to LPS. Similar effects we have previously observed in EA.hy926 cells stimulated with TNFα [Drabarek et al., 2012; Dymkowska et al., 2019]. Effect of LPS on metabolite profile changes over time of the treatment and indicates a reversible inhibition of pyruvate kinase, the enzyme that controls the intensity of PEP utilization for pyruvate formation. While the first phase of the changes (increased PEP level) does not have a clear explanation, progressive accumulation of pyruvate seems to be in line with LPS-induced elevation of PKM2 (pyruvate kinase isoform M2) activity that was observed in macrophages exposed to lipopolysaccharide for 24 h [Palsson-McDermott et al., 2015]. These effects are accompanied by substantially elevated levels of Krebs cycle intermediates probably potentiated due to inhibited mitochondrial NADH reoxidation through the respiratory chain. An increased level of lactate confirms a rise in anaerobic processes which allow cells to restore the pool of NAD^+^. Reduced oxidative metabolism and stimulation of glycolysis were previously described for various experimental models of inflammation [Robb et al., 2020; Vijayan et al., 2019; Yarbro et al., 2019]. Relatively stable or only slightly reduced level of ATP indicates that the cellular bioenergetics despite its shift towards a more glycolytic mode remains sufficient. On the other hand, some elevation in the AMP content might suggest activation of AMPK. The latter assumption is additionally strengthened as the proportion of phosphorylated form of ACC is increased in LPS-treated HAECs. A putative mechanism that is behind reduced oxidative phosphorylation seems not to be clear [Kelly and O’Neill, 2015]. Moderately increased cytosolic Ca^2+^ concentration should accelerate mitochondrial energy processes by stimulation of the TCA cycle. However, inhibition of the respiratory chain may have a superior effect and secondarily inhibits downstream metabolic processes delivering reduced nucleotides (NADH and FADH2) such as the TCA cycle. Thus glycolysis is a metabolic valve that allows cells to produce ATP in a mitochondria-independent manner and LPS makes endothelial cells more glycolytic than is observed in control conditions.

It cannot also be excluded that changes in the intensity of cellular calcium response are secondary and attributed to the reorganized mitochondrial network architecture and therefore affected the ability of mitochondria to buffer cytosolic Ca^2+^. The transient increase in ACC phosphorylation shown in Fig. 2 reflects an elevated AMPK activity and corresponds with the time course of changes in cellular lactate content. Interestingly, elevated activity of AMPK was implicated as one of the factors in a down-regulation of glycolysis in vascular endothelial cells subjected to pulsatile shear stress [Han et al., 2021]. However, in our stable experiments, shear stress was not considered. AMPK was found to indirectly induce PDK4 gene expression thereby inhibiting pyruvate dehydrogenase activity [Fritzen et al., 2015]. In contrast, AMPK also activates PDHc through phosphorylating catalytic subunit PDHA which alleviates inhibitory phosphorylation catalyzed by PDKa. It stimulates the TCA cycle as was observed in metastatic cancer cells [Cai et al., 2020] and is not in line with the data presented in this study. It seems that AMPK interacts with several and sometimes counteracting processes so the resultant, tissue-specific effect reflects precisely regulated and changing metabolic balance. It seems that the putative role of AMPK in the endothelial response to LPS is worthy of further study.

Cellular concentration of crucial metabolic intermediates has a key role in proper metabolic regulation and coordination of multiple metabolic processes and may switch metabolic processes from one mode to another. One of such factors is NAD^+^, a broadly used electron and proton acceptor in many intracellular redox reactions. Its reduced form (NADH) is oxidized in the respiratory chain which is a major energy transducing process in aerobic organisms (excluding photosynthesis in plants and other autotrophs). Moreover, NAD^+^ is used as a cofactor for sirtuins which are crucial metabolic regulators in mammals. Therefore, the cellular concentration of NAD^+^ must be controlled and maintained at an appropriate level. This in turn implies that excessive consumption of NAD^+^ precursors by alternative processes may be a limiting factor for NAD^+^ formation. Recently great attention has been focused on nicotinamide methylation catalyzed by nicotinamide N-methyltransferase (NNMT). This reaction competes for nicotinamide with nicotinamide phosphoribosyltransferase (NAMPT) and potentially may limit NAD^+^ synthesis. Moreover, nicotinamide methylation needs a methyl group donor that is S-adenosyl methionine (SAM). Therefore, an elevated NNMT activity may also affect other SAM-mediated methylation reactions. A growing body of evidence indicates that NNMT has a crucial regulatory function influencing not only energy metabolism but also epigenetic regulation in a variety of tissues [Sperber et al., 2015; Ulanovskaya et al., 2013; and for rev. Roberti et al., 2021]. However, the physiological role of this enzyme in vascular endothelial cells is still obscure. Experimental data concerning NNMT clearly show that cells challenged by some proinflammatory stimuli exhibit an elevated level of this protein[Dymkowska et al., 2021]. This may suggest that NNMT may play a role in the adaptive stress response of cells. On the other hand, its activation may contribute to the cellular harmful processes in stressful conditions. In fact, an elevated NNMT level was found in a course of some pathologies including hepatocellular carcinoma, breast cancer and stroma of colorectal cancer [Eckert et al., 2019; Li et al., 2019; Yang et al., 2021; Wang et al., 2019].

Here we have observed a time-dependent elevation of NNMT protein level in cells treated with LPS. However, we have not confirmed an expected elevation of MNA level (see Fig. 7, the first bars in each time point). In contrast, it was substantially reduced as soon as 30 min after the addition of LPS to the growth medium and though gradually increased together with increasing NNMT level, however, it never reached the control value found in cells untreated with LPS. It suggests additional effects of lipopolysaccharide attenuating an activity of NNMT-catalysed reaction. Moreover, we have not found any differences in NAD^+^ level in comparison to control cells regardless of the time of treatment with LPS (Fig. 8). Maybe NAD^+^ synthesis in HAECs occurs through alternative pathways, thus changes in nicotinamide level have not any impact on this process [Covarrubias et al., 2021; Liu et al., 2018].

### Silencing of NNMT-encoding gene prevents LPS-evoked effects

The direct link between LPS-evoked elevation of NNMT content and changes in the mitochondrial organization seems to be ambiguous. LPS only transiently affects mitochondrial network architecture which coincides with substantial changes in the NNMT level (see Fig. 4) and recoveries after prolonged treatment (see. Fig. 1). On the other hand, however, silencing of the NNMT encoding gene and therefore substantial reduction of NNMT protein level protects the mitochondrial network from LPS-induced reorganization (see Fig. 9). This discrepancy may suggest that the LPS-evoked effect on the NNMT level is not a direct cause of changes in the mitochondrial network distribution within cells but a reduction of NNMT level prior to stimulation of cells with LPS somehow prevents its detrimental effect. On a basis of the presented data, it is difficult to conclude the mechanism behind a counteracting effect of NNMT gene silencing on the metabolic activity of HAEC cells treated with LPS. Interestingly silencing of the NNMT gene itself does not influence the level of metabolites but fumarate whose concentration was substantially reduced. It could result from increased methylation of SDH gene promoter and reduced SDH transcription rate [Aggarwal et al., 2021, and for rev. Hawkins et al., 2018]. However, this assumption requires further experiments to test whether a decrease in the NNMT activity is accompanied by an increased s-adenosylmethionine concentration. Silencing of the NNMT gene tuned out to be very efficient (at least 60% reduction of the protein level) and reproducible. However, it needed a prolonged incubation with siRNA, thus we were conscious of the risks that it can activate cellular response prior to the addition of LPS.

To conclude, the aforementioned results are not sufficient to coherently explain the link between LPS-induced effects in HAECs and NNMT activity. One could speculate that aberrant Ca^2+^ homeostasis may be involved but still, it needs additional experimental confirmation. Recently a mechanism behind the activation of SOCE in HUVEC cells treated with LPS has been precisely described [Qiu et al. 2021]. Experiments shown here fully confirm those data showing substantially increased calcium response of HAEC cells exposed to LPS upon SOCE activation with thapsigargin. Silencing of the NNMT-encoding gene slightly counteracts such a response, which makes the role of Ca^2+^ signalling in this context unclear potentially relevant. It is worthy of notice that all effects of LPS we have observed are substantially prevented if the NNMT-encoding gene is silenced. We suppose that restoration of calcium signalling is at the bottom of the remaining effects of NNMT elimination. However, this tempting speculation needs deeper study to be confirmed.

It seems to be clear that silencing of the NNMT encoding gene and therefore reduction of the NNMT protein content and activity influences HAECs in a metabolic point(s) located “above” several important regulatory phenomena such as AMPK activation, calcium response and rearrangement of mitochondrial network architecture and some other not shown here as exceeding the scope of the presented study. Thus, regulation of NNMT activity is of paramount importance for crucial metabolic processes in endothelial cells, which outreach earlier accepted the vasculoprotective role of this enzyme by delivering MNA [Fedorowicz et al., 2016; Przyborowski et al., 2015]. Because of the fact that NNMT stands at the crossroads of metabolic pathways and epigenetic regulatory processes the complete understanding of its role not only in the endothelial cells but also in many others seems to be very difficult and requires multilateral approaches. It is tempting to speculate that NNMT could be considered a potential therapeutic target and this aspect was also pointed out by others [Gao et al., 2021]. However, the manifold functions of this enzyme must be carefully counted.

## Acknowledgements

We thank Prof. Stefan Chłopicki from JCET (Jagiellonian University, Cracow) for inspiring discussions and our colleague Dr. Adam Jagielski from the Institute of Biochemistry, Faculty of Biology at the University of Warsaw for assistance in metabolite measurements.

